# Maternal psychosocial risk factors and offspring gestational epigenetic age acceleration in a South African birth cohort study

**DOI:** 10.1101/503623

**Authors:** Nastassja Koen, Meaghan Jones, Raymond T. Nhapi, Kirsten A. Donald, Whitney Barnett, Nadia Hoffman, Julia L. MacIsaac, Alexander M. Morin, David T.S. Lin, Michael S. Kobor, Karestan C. Koenen, Heather J. Zar, Dan J. Stein

## Abstract

Epigenetic age (EA) acceleration is associated with higher risk of chronic disease and mortality in adults. However, little is known about whether and how *in utero* exposures might shape gestational EA acceleration at birth. We aimed to explore associations between maternal psychosocial risk factors and offspring gestational EA acceleration at birth in a South African birth cohort study – the Drakenstein Child Health Study. Maternal psychosocial risk factors included trauma/stressor exposure; posttraumatic stress disorder (PTSD); depression, psychological distress; and alcohol/tobacco use. Offspring gestational EA acceleration at birth was calculated using an epigenetic clock previously devised for neonates. Bivariate linear regression was used to explore unadjusted associations between maternal risk factors and offspring gestational EA acceleration at birth. A stepwise regression method was then used to determine the best multivariable model for adjusted associations. Data from 272 maternal-offspring dyads were included in the current analysis. In the stepwise regression model, maternal trauma exposure (*β*=7.92; *p*<0.01) or PTSD (*β*=7.46; *p*<0.01) were significantly associated with offspring gestational EA acceleration at birth, controlling for ethnicity, offspring sex, head circumference at birth, maternal HIV status, and prenatal tobacco or alcohol use. In site-stratified models, these associations retained statistical significance and direction of effect. Maternal trauma exposure or PTSD may thus be associated with offspring gestational EA acceleration at birth. Given the novelty of this preliminary finding, and its potential translational relevance, further studies to delineate underlying biological pathways and to explore clinical implications of EA acceleration are warranted.

## INTRODUCTION

Maternal exposure to psychosocial risk factors during pregnancy – including traumatic stressors, psychiatric disorders and symptoms, and substance misuse – constitute a notable public health concern; and may be associated with poor birth outcomes and adverse health and development in affected offspring^1,2^. Given the high burden of psychosocial risk factors in low- and middle-income countries (LMICs) such as South Africa^3^, maternal-offspring dyads in these settings are particularly vulnerable. Prenatal psychosocial risk exposure may be associated with measurable changes in DNA methylation (DNAm) at specific genomic sites in the affected offspring^4^. It is noteworthy, though, that a large meta-analysis found no large epigenome-wide effects of prenatal maternal stress exposure on offspring differential DNAm^5^. Recent work has also investigated DNAm as a predictor of chronological age in children and adults^6,7^. However, given the limitations in transferability of such ‘epigenetic clocks’ (which were calibrated initially for adult populations); specialised clocks have recently been generated to estimate gestational age at birth^8-10^.

Epigenetic age (EA) acceleration – greater DNAm gestational age relative to chronological age – has been found to be associated with increased risk of age-related disorders and mortality in adults^11,12^. Further, recent work has reported significant associations between a number of maternal-offspring biomedical factors (eg. pre-pregnancy maternal obesity; maternal age over 40 years at delivery; lower offspring birthweight; lower 1-minute Apgar score) and EA acceleration at birth in the affected offspring^13,14^. However, EA deceleration (ie. lower epigenetic age relative to chronological age) associated with maternal risk (eg. insulin-treated gestational diabetes mellitus) has also been reported^13^. Thus, the functional implications of epigenetic age acceleration/deceleration remain unclear^13,15^. There is also a paucity of existing literature exploring maternal psychosocial predictors of offspring gestational EA acceleration, particularly in LMICs.

This analysis was nested within the Drakenstein Child Health Study (DCHS), an ongoing, interdisciplinary, longitudinal birth cohort study investigating determinants of maternal-child health in a poor, peri-urban South African community^16-18^. We have previously reported a high burden of maternal psychosocial risk factors (eg. traumatic stress exposure/posttraumatic stress disorder (PTSD), alcohol and tobacco use); and significant associations between these factors and adverse offspring birth and neurodevelopmental outcomes in this cohort^19-23^. In the current analysis, we aimed to investigate associations between maternal psychosocial risk factors and offspring EA acceleration at birth. We hypothesised that key maternal risk factors would be significantly associated with offspring gestational EA acceleration in this study sample.

## MATERIALS/SUBJECTS AND METHODS

### Participants

Pregnant women were recruited at 20-28 weeks’ gestation from two primary care clinics in the Drakenstein sub-district in Paarl, Western Cape, South Africa – one (TC Newman) serves a predominantly Mixed ancestry community; while the other (Mbekweni) serves primarily a Black/African ancestry community. Within the DCHS, mothers are followed throughout pregnancy and childbirth until the index child is at least 5 years old^16-18^. General inclusion and exclusion criteria for the DCHS are described fully elsewhere^17^.

For this analysis, only samples of mothers with complete and accurate sociodemographic and psychosocial phenotype data; with recorded maternal informed consent for the collection, storage, and future analyses of offspring DNA; and with non-contaminated offspring Illumina Infinium HumanMethylation450K or MethylationEPIC BeadChip data were eligible for inclusion^24^ (see also *Statistical Analyses*, below).

This study sample included mothers enrolled into the larger study between March 2012 and March 2015. From a total of 1225 mothers enrolled, 1143 were retained in the cohort at the time of birth; and delivered live births. From this sub-sample, 276 mothers had offspring with non-contaminated HumanMethylation450K (n=120) or MethylationEPIC (n=156) BeadChip data; and complete maternal/offspring phenotype data. This sub-sample was initially selected for analysis based on a number of risk criteria of relevance to the larger DCHS – including (but not limited to) maternal exposure to psychological trauma and/or PTSD. Two sets of twins was excluded, due to concerns of analytic mixed effects. Thus, data from a sub-sample of 272 maternal-offspring dyads were included in the current analysis.

### Variables and Measurement

Maternal sociodemographic and antenatal psychosocial risk factors; offspring anthropometric outcomes; and offspring gestational EA acceleration at birth were assessed, as is detailed below. Maternal psychosocial assessment in the DCHS has been described fully previously^17^.

#### Maternal sociodemographic characteristics

Socioeconomic status (SES) and related sociodemographic characteristics were assessed using a questionnaire adapted from the South Africa Stress and Health Study (SASH)^25^.

#### Maternal antenatal psychosocial risk factors

Exposure to stressful events – either recently or in one’s lifetime – was assessed using the modified World Mental Health Life Events Questionnaire (LEQ; adapted for our purposes from the SASH^25^); the Childhood Trauma Questionnaire (CTQ)^26^; and the Intimate Partner Violence (IPV) Questionnaire (adapted from the WHO Multicountry Study^27^ and the Women’s Health Study (Zimbabwe))^28^. The Modified PTSD Symptom Scale (MPSS)^29^ was used as a rapid screening for psychological trauma exposure and/or PTSD, based on the DSM diagnostic criteria and adapted for our purposes to incorporate a more “broad” approach to defining the index traumatic events (eg. see Stein et al. 2014^30^ for “narrow” vs “broad” approaches for diagnosing PTSD). The MPSS screens for PTSD symptoms in the preceding 2 weeks. Psychological distress was assessed using the Self-Reporting Questionnaire (SRQ-20)^31,32^; and depression using the Beck Depression Inventory II (BDI-II)^33-35^ and the Edinburgh Postnatal Depression Scale (EPDS)^36,37^. Maternal substance misuse was identified using the World Health Organization’s (WHO’s) Alcohol, Smoking and Substance Involvement Screening Test (ASSIST)^38-40^ and a retrospective Alcohol Exposure Questionnaire (AEQ). The AEQ was designed for the purposes of the DCHS to quantify alcohol use (during any of the three trimesters of pregnancy) in mothers assessed as high-risk on the ASSIST.

#### Offspring anthropometry at birth

Offspring weight, head circumference, and height/length at birth were measured by trained clinical staff, and the relevant z scores were then calculated using the Fenton preterm growth charts^41,42^. Following the WHO’s convention, low weight-for-age z score (WAZ), low head circumference-for-age z score (HCAZ) and low height-for-age z score (HAZ) each was defined as a score of 2 standard deviations or more below the mean^43^. Prematurity was defined as birth before 37 completed weeks’ gestation; and low birthweight as less than 2,500g.

#### Umbilical cord blood collection

In the DCHS, umbilical cord blood was collected by trained staff after offspring delivery but before delivery of the placenta. The cord was clamped and cut, after which the clamp was released and cord blood drained by gravity into a kidney dish. Thereafter, cord blood was collected using a syringe for processing and storage^24^. Staff were specifically trained to ensure only cord blood drained into the collection dish and that to their best ability, no other/external blood was collected.

### Ethical Considerations

The DCHS was approved by the Human Research Ethics Committee (HREC) of the Faculty of Health Sciences, University of Cape Town (UCT) and by the Western Cape Provincial Research Committee. Within the DCHS, all potential participants were provided with a consent form describing the scope and aims of the study, in their preferred language (English, Afrikaans or isiXhosa). Following verbal and written informed consent, mothers were asked to complete a battery of self-report measures at an antenatal study visit (28-32 weeks’ gestation). While maternal phenotype data from a number of postnatal timepoints are also collected within the larger DCHS^16,17^; the current analysis pertains to the antenatal maternal assessment only.

All measures were administered by trained study fieldworkers in either English, Afrikaans or isiXhosa, as per participants’ preferences. Interviews were conducted in private onsite consultation rooms and every effort was made by study staff to ensure confidentiality. Participants were also provided with refreshments and standard reimbursement for transport costs. On completion of the study assessment, those participants with suspected psychopathology or psychosocial risk (eg. trauma/IPV exposure, PTSD, depression and/or substance use) were referred by study staff to the most appropriate care providers in the community, according to a standard operating procedure devised for the purposes of the DCHS. Health promotion information leaflets designed by the study team (and translated into participants’ preferred language) were also offered to all enrolled mothers, and included contact details for local and accessible health service providers.

### Statistical Methods

Data were analysed using R^44^ and STATA^45^.

#### Maternal-offspring phenotype

Frequency distributions and medians (interquartile ranges) were used to describe maternal sociodemographic characteristics and psychosocial risk profile; as well as infant anthropometric parameters at birth.

#### Offspring epigenetic age acceleration

As has been described fully previously^24^, offspring DNAm was measured using the Illumina Infinium HumanMethylation450K (Illumina, San Diego, USA) or the MethylationEPIC BeadChip^46^ as per manufacturers’ instructions. Raw data were imported into Illumina GenomeStudio Software for background subtraction and colour correction, and then exported for processing using the *lumi* package in R (version 3.2.3). Quality control procedures were standardised and are discussed in detail elsewhere^24^.

Offspring gestational EA acceleration at birth was calculated using an epigenetic clock particularly suited for newborns which outputs gestational age^8,10^ and is based on prior work in adult populations using chronological age^6^. This method employs a panel of CpGs to predict chronological age, of which 8 are missing on the EPIC array. For EPIC data, these 8 CpGs were included with values set to NA. In the current analysis, gestational EA acceleration was calculated as the residuals of the linear model between epigenetic gestational age and chronological GA^47^.

#### Associations between maternal psychosocial risk factors and offspring gestational epigenetic age acceleration at birth

Bivariate analyses were used to explore unadjusted associations between key maternal sociodemographic and psychosocial risk variables (eg. income, trauma/stress, PTSD, psychological distress, depression, tobacco/alcohol use and maternal HIV infection) and offspring gestational EA acceleration at birth. Thereafter, a stepwise model selection method was used to obtain the best multiple regression model (from which the adjusted associations were obtained), based on minimizing the Akaike Information Criterion (AIC). Likelihood ratio tests were then used to eliminate any statistically insignificant variables from this model. Given prior site-specific differences in key exposure variables such as tobacco/alcohol use^17^ and maternal HIV (see below), the regression analysis was repeated, stratified by study site.

## RESULTS

### Maternal sociodemographic characteristics

This study sample comprised 151 (56%) women recruited from Mbekweni and 121 (44%) from TC Newman. At enrolment, the median (IQR) age of mothers was approximately 26 (22; 31) years, **Table 1**. Approximately a third were primigravid, and most (61%) were unmarried/not co-habiting with their partner. Despite approximately half of the study sample having completed some secondary education, nearly three quarters (73%) were unemployed and the vast majority (84%) reported a monthly household income of ≤ ZAR 5000 (≈USD 400). While the prevalence of maternal HIV infection in the study sample was 24%, a statistically significant site-specific difference was noted (p<0.01), with 6% of mothers at TC Newman HIV infected, versus 38% at Mbekweni. No offspring in this study sample was HIV infected.

**TABLE 1:**
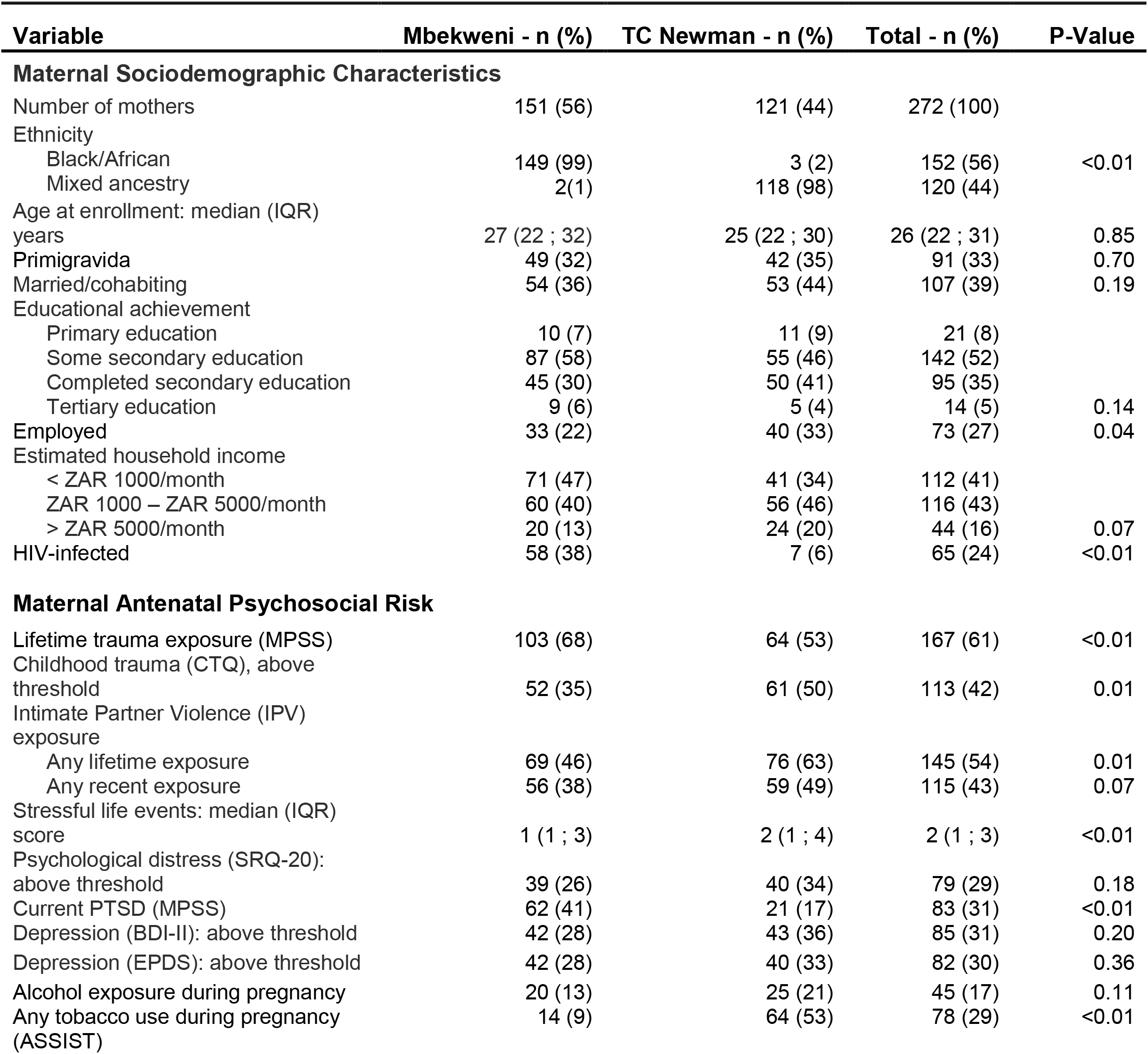
Maternal sociodemographic characteristics and antenatal psychosocial risk

### Maternal antenatal psychosocial risk

Most mothers (61%) in this study sample reported exposure to any psychological trauma during their lifetimes; 42% had been exposed to trauma (abuse/neglect) during childhood; and the prevalence of lifetime exposure to intimate partner violence (IPV) was 54% (with 43% having experienced IPV during the past year), **Table 1**. The prevalence of current PTSD was 31%. Despite a relatively low median (IQR) score on the modified Life Events Questionnaire [2 (1; 3)], 29% of the study sample reporting symptoms of psychological distress on the SRQ-20. Similarly, almost a third scored above threshold for depression on both the BDI-II (31%) and the EPDS (30%). Alcohol consumption during pregnancy was reported in 17% of participants, and prenatal tobacco use in 29%. A notable and statistically significant site-specific difference was evident in the latter, with 9% of mothers from Mbekweni and 53% from TC Newman having smoked tobacco during pregnancy (p<0.01).

### Offspring anthropometry at birth

More than half (56%) of offspring in this study sample was male; the median (IQR) gestational age at birth of 39 (38; 40) weeks, **Table 2**. Almost a fifth of offspring (19%) were born pre-term; 11% had low birth weight; 8% had decreased weight-for-age z (WAZ) scores at birth and 12% had low head-circumference-for-age z (HCAZ) scores. The median (IQR) weight at birth in this study sample was 3.1 (2.8; 3.5) kg; and the median (IQR) head circumference was 34 (33; 35) cm.

**TABLE 2:**
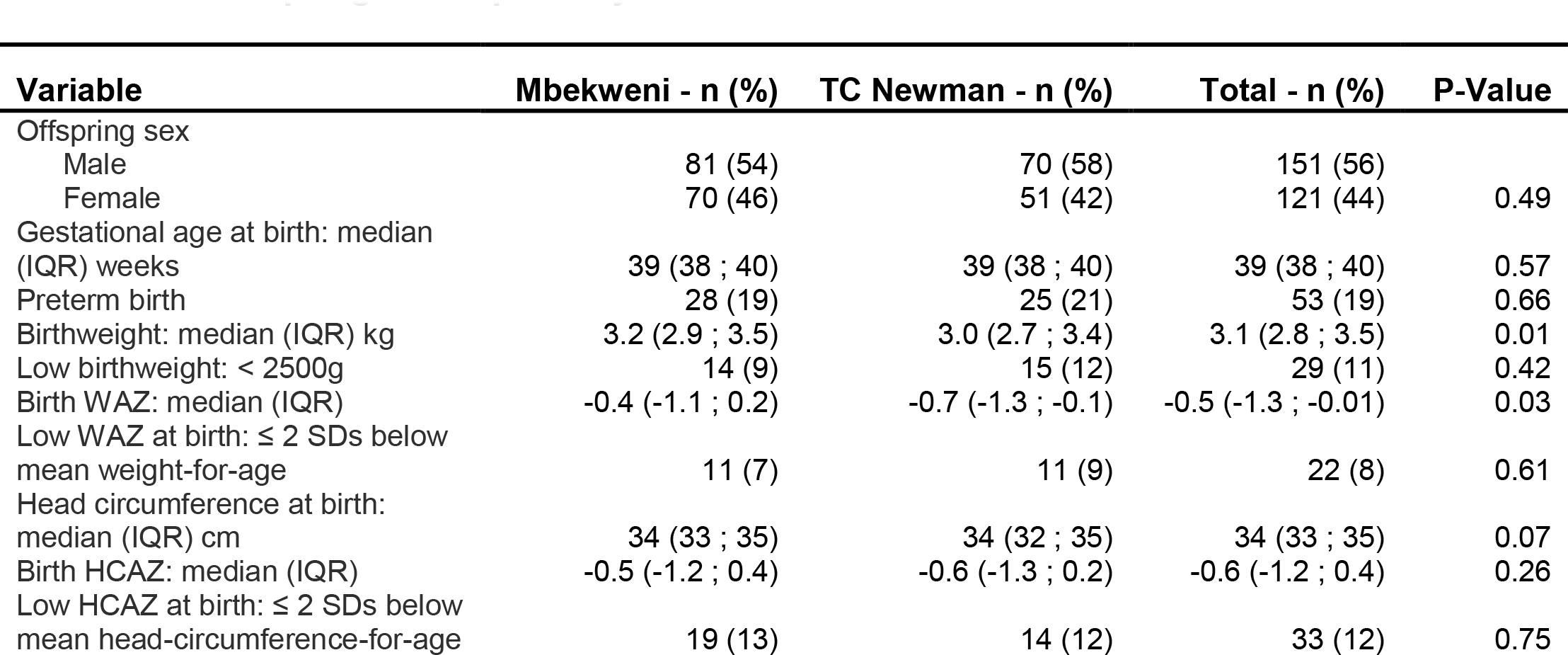
Offspring anthropometry at birth

### Associations between maternal psychosocial risk and offspring gestational epigenetic age acceleration at birth

In bivariate unadjusted analyses, ethnicity (*β*=-5.53; *p*<0.01), study site (β=-5.23; p<0.01), offspring sex (*β*=-3.36; *p*=0.02), head circumference at birth (*β*=0.82; *p*=0.03), maternal trauma exposure (*β*=8.03; *p*<0.01) or PTSD (*β*=9.25; *p*<0.01) each was found to be significantly associated with offspring gestational EA acceleration at birth. In the final stepwise regression model, maternal trauma exposure (*β*=7.92; *p*<0.01) or PTSD (*β*=746; *p*<0.01) each was significantly associated with offspring gestational EA acceleration at birth (ie. the residuals of the linear model between scaled epigenetic gestational age and chronological gestational age), when controlling for ethnicity, offspring sex, head circumference at birth, maternal HIV status, and prenatal tobacco or alcohol use, **Table 3**. When stratifying the model by study site, these associations retained statistical significance and direction of effect, **Table 3**. When employing a more “narrow” definition of trauma exposure/PTSD - which adheres strictly to the DSM-5 criteria^48^ - maternal trauma exposure, but not PTSD, was found to be significantly associated with offspring gestational EA acceleration at birth in the full study sample (*β*=5.54; *p*<0.01) and the TC Newman subset (*β*=6.04; *p*=0.01); and tending towards significance in the Mbekweni subset (*β*=4.94; *p*=0.06).

**TABLE 3:** Associations between maternal psychosocial risk factors and offspring gestational epigenetic age acceleration at birth

|  | Unadjusted<br>β Estimate [95% CI] | P-Value | Adjusted<br>β Estimate [95% CI] | P-Value |
| --- | --- | --- | --- | --- |
| **TOTAL STUDY SAMPLE** |  |  |  |  |
| Ethnicity: Mixed ancestry | -5.53 [-8.24; -2.82] | <0.01** | -5.79 [-9.13; -2.45] | <0.01** |
| Study site: TC Newman | -5.23 [-7.95; -2.52] | <0.01** | N/A | N/A |
| Offspring sex: Male | -3.36 [-6.12; -0.61] | 0.02** | -3.67 [-6.32; -1.03] | 0.01** |
| Offspring head circumference at birth | 0.82 [0.06; 1.58] | 0.03** | 0.70 [-0.03; 1.42] | 0.06 |
| Maternal HIV status: Positive | -0.70 [-3.95; 2.55] | 0.67 | -3.09 [-6.38; 0.19] | 0.07 |
| Alcohol exposure during pregnancy | -1.20 [-4.94; 2.53] | 0.53 | -0.21 [-3.95; 3.53] | 0.91 |
| Tobacco use during pregnancy | -1.56 [-4.62; 1.51] | 0.32 | 2.76 [-0.73; 6.25] | 0.12 |
| Maternal trauma exposure/PTSD |  |  |  |  |
| Trauma exposure | 8.03 [4.85; 11.22] | <0.01** | 7.92 [4.80; 11.04] | <0.01** |
| PTSD | 9.25 [6.06; 12.44] | <0.01** | 7.46 [4.13; 10.80] | <0.01** |
| **MBEKWENI** |  |  |  |  |
| Offspring sex: Male | -4.20 [-7.97; -0.42] | 0.03** | -5.24 [-9.10; -1.37] | <0.01** |
| Offspring head circumference at birth | 1.28 [0.15; 2.41] | 0.03** | 1.34 [0.18; 2.49] | 0.02** |
| Maternal HIV status: Positive | -3.38 [-7.27; 0.52] | 0.09 | -2.20 [-6.03; 1.64] | 0.26 |
| Alcohol exposure during pregnancy | -0.98 [-6.66; 4.70] | 0.73 | 0.56 [-5.64; 6.75] | 0.86 |
| Tobacco use during pregnancy | 0.60 [-6.03; 7.24] | 0.856 | 0.65 [-6.46; 7.76] | 0.86 |
| Maternal trauma exposure/PTSD |  |  |  |  |
| Trauma exposure | 8.22 [3.28; 13.15] | <0.01** | 8.05 [3.11; 12.99] | <0.01** |
| PTSD | 9.43 [4.94; 13.92] | <0.01** | 8.36 [3.78; 12.95] | <0.01** |
| **TC NEWMAN** |  |  |  |  |
| Offspring sex: Male | -1.81 [-5.67; 2.05] | 0.35 | -2.08 [-5.81; 1.65] | 0.27 |
| Offspring head circumference at birth | 0.09 [-0.89; 1.06] | 0.86 | 0.31 [-0.64; 1.25] | 0.52 |
| Maternal HIV status: Positive | -4.37 [-12.51; 3.78] | 0.29 | -6.42 [-14.51; 1.66] | 0.12 |
| Alcohol exposure during pregnancy | -0.16 [-4.88; 4.56] | 0.95 | -0.24 [-5.03; 4.55] | 0.92 |
| Tobacco use during pregnancy | 1.87 [-1.94; 5.68] | 0.33 | 2.72 [-1.19; 6.64] | 0.17 |
| Maternal trauma exposure/PTSD |  |  |  |  |
| Trauma exposure | 7.56 [3.54; 11.58] | <0.01** | 7.95 [3.93; 11.97] | <0.01** |
| PTSD | 5.66 [0.58; 10.74] | 0.03** | 6.72 [1.35; 12.09] | 0.02** |

\*\*p<0.05

## DISCUSSION

In this study of the association between maternal exposure to prenatal psychosocial risk factors and gestational epigenetic age (EA) acceleration at birth in the index offspring, we found that infants born to mothers with trauma exposure or PTSD had gestational EA acceleration versus those without such exposure, when controlling for a number of relevant covariates.

These findings are in line with emerging evidence of trauma-/PTSD-associated EA acceleration in adult combat veterans^49,50^. One recent meta-analysis by the Psychiatric Genomics Consortium PTSD Epigenetics Workgroup across 9 adult cohorts (combined N=2186, including seven military samples) also reported small but significant associations each between childhood trauma exposure and severity of lifetime PTSD, and EA acceleration^51^. However, neither PTSD diagnosis nor lifetime trauma exposure was associated with EA acceleration in this meta-analysis; and a small longitudinal study of male soldiers (n=96), assessing pre- and post-deployment, reported an inverse association between the development of PTSD symptoms and EA acceleration (although trauma was found to be significantly associated with EA acceleration in this study)^52^. Further, while a small recent study (N=101) reported EA acceleration in children exposed directly to neighbourhood violence^53^; EA deceleration has also been found to be associated with higher offspring distress in the context of low caregiver contact^15^. Thus, it may be that offspring EA acceleration or deceleration signifies exposure to prenatal maternal risk^13^.

A number of limitations of the current study should be acknowledged. First, the sample size was relatively small, thus decreasing the statistical power of association analyses. Nevertheless, we report important associations with trauma or PTSD exposure. Second, self-report questionnaires to assess maternal exposure to psychosocial risk factors may have resulted in under-reporting of key variables such as tobacco and/or alcohol use. Given prior evidence of associations between maternal smoking/alcohol consumption and deviations in EA^54^, self-reporting bias may have contributed to the lack of significant associations in our analysis. However, we have previously shown that self-reported tobacco smoking correlated well with urine cotinine measurement (an objective biomarker of tobacco smoking/exposure) in our study cohort; particularly in the Mixed ancestry sample^23^. Third, PTSD was not assessed using a clinician-administered diagnostic interview, which may have biased the findings.

These limitations notwithstanding, our study provides a unique preliminary exploration of associations between maternal psychosocial risk factors and offspring gestational EA acceleration at birth. To the best of our knowledge, an association between maternal trauma exposure/PTSD and gestational EA acceleration in the index offspring has not previously been reported. Future studies – incorporating larger samples sizes and/or cross-tissue analyses – are warranted in order to explore such associations further; and to delineate underlying neurobiological mechanisms. In addition, work is needed to understand the implications of EA acceleration at birth. For example, it is currently unknown whether EA acceleration is also associated with adverse neurodevelopmental outcomes in the affected offspring. Further, while the adverse intergenerational effects of maternal trauma exposure and PTSD have been well documented – both in the DCHS cohort^19–21^ and globally^1,55^ – the potential role of EA acceleration as a driver of these associations is yet to be determined. From a translational perspective, if EA acceleration at birth does indeed prove clinically meaningful in future, it may contribute to the development of trauma-/PTSD-specific epigenetic biomarkers to inform maternal-offspring screening, diagnostic and therapeutic interventions.

## ACKNOWLEDGEMENTS

This study was supported by the Eunice Kennedy Shriver National Institute of Child Health and Human Development of the National Institutes of Health (NICHD) under Award Number R21HD085849, the Fogarty International Center (FIC) and the Bill and Melinda Gates Foundation (OPP 1017641). The content is solely the responsibility of the authors and does not necessarily represent the official views of the National Institutes of Health.

Additional support for HJZ, DJS, NK and WB, and for research reported in this publication was by the South African Medical Research Council (SAMRC); from a Newton Advanced Fellowship (KD); and from South Africa’s National Research Foundation (NRF) (HZ; grant number: 105865). WB is supported by the SAMRC National Health Scholars programme; NK and KD receive additional support from the SAMRC under Self-Initiated Research Grants. The views and opinions expressed are those of the authors and do not necessarily represent the official views of the SAMRC.

We thank the Drakenstein Child Health Study staff, and the clinical and administrative staff of the Western Cape Government Department of Health at Paarl Hospital and at the clinics for support of the study. We also thank our collaborators and students. Finally, we thank all mothers and children enrolled in the Drakenstein Child Health Study.

